# Mitotic progression following DNA damage enables pattern recognition within micronuclei

**DOI:** 10.1101/156414

**Authors:** Shane M Harding, Joseph L Benci, Jerome Irianto, Dennis E Discher, Andy J Minn, Roger A Greenberg

## Abstract

Inflammatory gene expression following genotoxic cancer therapy is well documented, yet the events underlying its induction remain poorly understood. Inflammatory cytokines modify the tumor microenvironment by recruiting immune cells and are critical for both local and systemic (abscopal) tumor responses to radiotherapy^1^. An enigmatic feature of this phenomenon is its delayed onset (days), in contrast to the acute DNA damage responses that occur in minutes to hours. Such dichotomous kinetics implicate additional rate limiting steps that are essential for DNA-damage induced inflammation. Here, we show that cell cycle progression through mitosis following DNA double-strand breaks (DSBs) leads to the formation of micronuclei, which precede activation of inflammatory signaling and are a repository for the pattern recognition receptor cGAS. Inhibiting progression through mitosis or loss of pattern recognition by cGAS-STING impaired interferon signaling and prevented the regression of abscopal tumors in the context of ionizing radiation and immune checkpoint blockade *in vivo*. These findings implicate temporal modulation of the cell cycle as an important consideration in the context of therapeutic strategies that combine genotoxic agents with immune checkpoint blockade.

---

Radiotherapy and many chemotherapeutics rely on DNA double strand break (DSB) formation to drive the killing of tumor cells over several cell division cycles^2,3^. Concomitant with this protracted cell death schedule, inflammatory cytokine production increases over days following the insult. As a host of cytokines and inflammatory signals are produced following ionizing radiation (IR)^4,5^, we used STAT1 phosphorylation at Y701 as a surrogate for inflammatory pathway activation (Fig. 1a and 1b). MCF10A mammary epithelial cells showed STAT1 activation between 3 and 6 days post-IR with a dose-dependent threshold of at least 5Gy. Total STAT1 protein and the mRNA levels of multiple inflammatory genes were also detected in a time dependent manner (Fig. 1c. and Extended data 1a). Using inducible nucleases (Fig. 1d and Extended data 1b and 1c), we observed a delayed accumulation of active STAT1 and inflammatory gene expression, confirming these signals are driven by DSBs.

**Figure 1:**
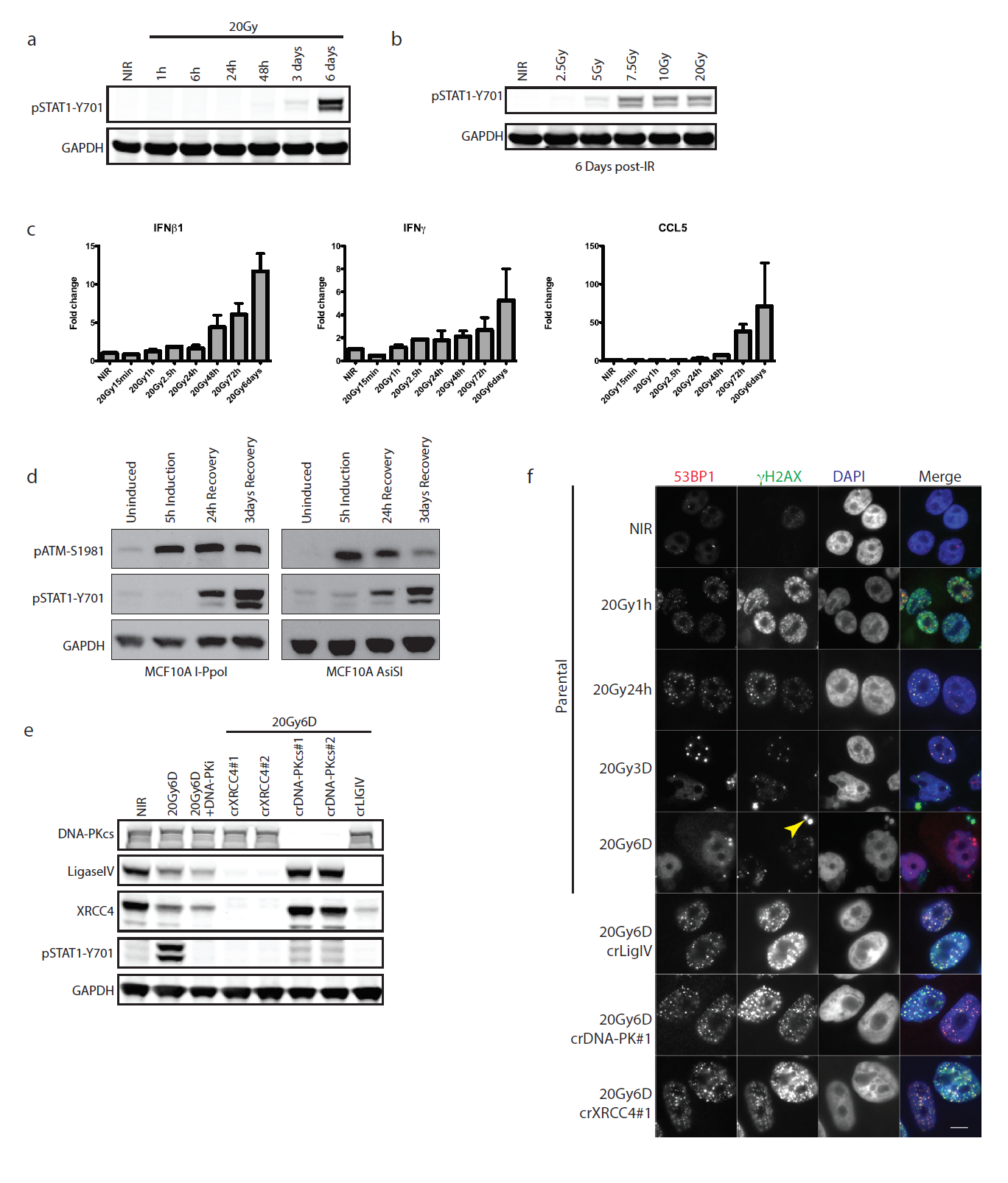
Loss of NHEJ antagonizes DSB-induced inflammatory signaling. **a,** Representative western blots for activation of STAT1 signaling at indicated times post-20Gy. NIR, non-irradiated. **b,** Dose-dependency of STAT1 response at 6 days as in panel (**a)**. **c,** RT-qPCR of inflammatory gene induction following IR. Error bars represent SEM of at least 3 biological replicates. **d,** STAT1 activation monitored as in (**a)**. Western blotting for STAT1 activation after 5h induction of DSBs by AsiSI or I-PpoI nuclease with Shield/4-OHT and followed by washout and recovery for the indicated times. **e,** CRISPR-Cas9 knockout of NHEJ components were monitored for STAT1 activation as in (**a)**. **f,** Immunofluorescence of 53BP1 and γH2AX indicating residual DSBs in crNHEJ cells. Arrowhead indicates γH2AX positive micronucleus. Scale bar is 10µm.

We reasoned that if residual DSBs were driving inflammatory signals then failure of non-homologous end-joining (NHEJ) DSB repair should amplify the response. Paradoxically, we observed that inhibition of DNA-PKcs (DNA-PKi) or CRISPR-Cas9 knockout of multiple NHEJ components diminished STAT1 activation in MCF10A and prostate epithelial cells (Fig 1e and Extended data 1c and 1d). STAT1 activation through exogenous IFNβ1 was unaffected by DNA-PKi (Extended data 1e), ruling out a direct role in STAT1 phosphorylation. Inhibition of ATM kinase on the other hand had little influence over the level of STAT1 activation (Extended data 1f).

DSB-induced STAT1 activation correlated with the appearance of aberrantly shaped nuclei and micronuclei (Fig 1f and 2a). The DSB marker γH2AX was increased in micronuclei however 53BP1 was absent, consistent with their reported defects in DSB signaling (Fig 1f)^6,7^. Nuclei of NHEJ knockout and DNA-PKi treated cells were morphologically normal despite being replete with DSBs (Fig 1f and 2a). As micronuclei are byproducts of mitotic progression, these data suggest that STAT1 activation occurred following mitosis^8^. Flow cytometry showed progression of parental cells from G2 into G1 between 24 and 48h post-IR, whereas NHEJ knockouts were static over this time (Extended data 2a). This corresponds to a wave of parental cells moving into mitosis as evidenced by phospho-histone H3 staining that is absent in the NHEJ knockout cells (Fig 2b). Notably, the specific CDK1 inhibitor RO-3306 (CDK1i) or PLK1 inhibitor BI-2536 (PLK1i) blocked mitotic entry, micronuclei formation and STAT1 activation (Figs 2a-2c and extended data 2b.)^9^. Furthermore inhibition of cell cycle progression by other CDKi reduced the flux of cells through mitosis and STAT1 activation (Extended data 2c). CDK1i also prevented STAT1 activation in PARPi treated BRCA1 mutated ovarian cancer cells (Fig 2d), suggesting mitotic entry as a gateway to inflammatory signaling from diverse DSB inducing stimuli.

**Figure 2:**
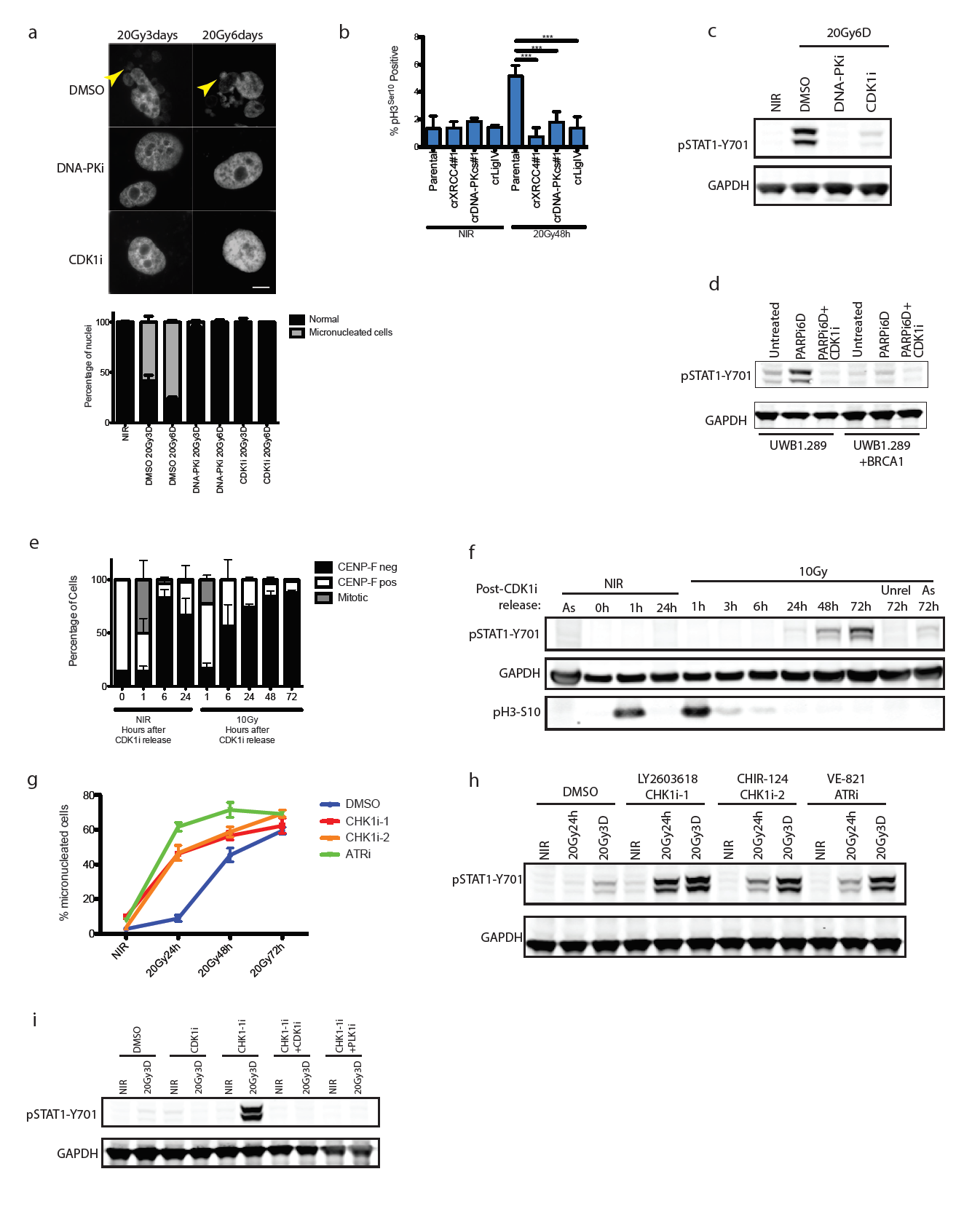
Progression through mitosis underlies inflammatory pathway activation. **a,** Representative images of DAPI-stained MCF10A nuclei. Quantification was as described in methods and error bars represent SEM. Arrowhead highlights micronuclei. Scale bar is 10µm. **b,** The mitotic fraction of cells was quantified using H3 (Ser10) phosphorylation measured by flow cytometry and expressed as a percentage of total single cells. ***p<0.0001 with Dunnett’s multiple comparison. **c,** Western blot of IR-induced STAT1 activation under the indicated conditions in MCF10A cells. **d,** Western blot of STAT1 activation in UWB1.289 or UWB1.289+BRCA1 reconstituted cells treated for 6 days with PARPi treatment with or without CDK1i. **e,** Cells were synchronized in G2 with CDK1i prior to irradiation and released for the indicated times. CENP-F positive (S/G2), CENP-F negative (G0/G1) and mitotic cells were quantified in 3 independent experiments. **f,** Representative western blot for STAT1 activation following the release scheme described in (**e).** As, (asynchronous). **g,** DAPI stained MCF10A cells were quantified as in (**a).** Following IR in the presence of the indicated inhibitors. Error bars represent SEM of three independent experiments. **h,** Representative western blot of STAT1 activation of cells treated as in **g. i,** Representative western blots of STAT1 activation with concurrent CHK1i and mitotic blockade.

To further test the requirement for mitotic progression, we synchronized cells in G2 before irradiation. Release from CDK1i after IR resulted in progression from G2 into the following G1 over 72h as determined by loss of the S/G2-phase and mitosis marker CENP-F (Fig 2e). STAT1 phosphorylation levels were greater than that observed in asynchronous cells and absent in cells not released from G2 (Fig 2f). The progression from G2 into mitosis after DSBs is biphasically controlled by an ATM dependent transient G2/M checkpoint and a slower G2 accumulation that requires ATR and Chk1 kinase activity^10-13^. CHK1 or ATR inhibition accelerated production of micronucleated cells, STAT1 activation and progression into G1 following damage (Fig 2g and 2h, and Extended data 2d). This accelerated response was completely abolished by concomitant CDK1i or PLKi (Fig 2i), emphasizing the need for passage through mitosis to drive DSB induced inflammatory signaling.

STAT1 phosphorylation can be elicited following activation of the DNA dependent pattern recognition receptor (PRR) cGAS and its downstream mediators STING and IRF3, which serve to transcriptionally activate type I interferon^14-22^. CRISPR-Cas9 knockout of either cytosolic cGAS or STING reduced activation of downstream IRF3 target genes ISG54, ISG56 and the NF-κβ target CCL5 post-IR (Fig 3a and Extended data 3a)^23,24^. Attenuated STAT1 activation and IFITM1 induction was also evident in cGAS-STING knockouts (Fig 3b and Extended data 3c). Senescence associated β-galatosidase staining was relatively unaffected by treatments that influence inflammatory activation (Extended data 3c), suggesting that alterations in epithelial cell senescence were not contributory to the reduced interferon signaling observed upon inhibiting mitotic entry or PRR activation.

**Figure 3:**
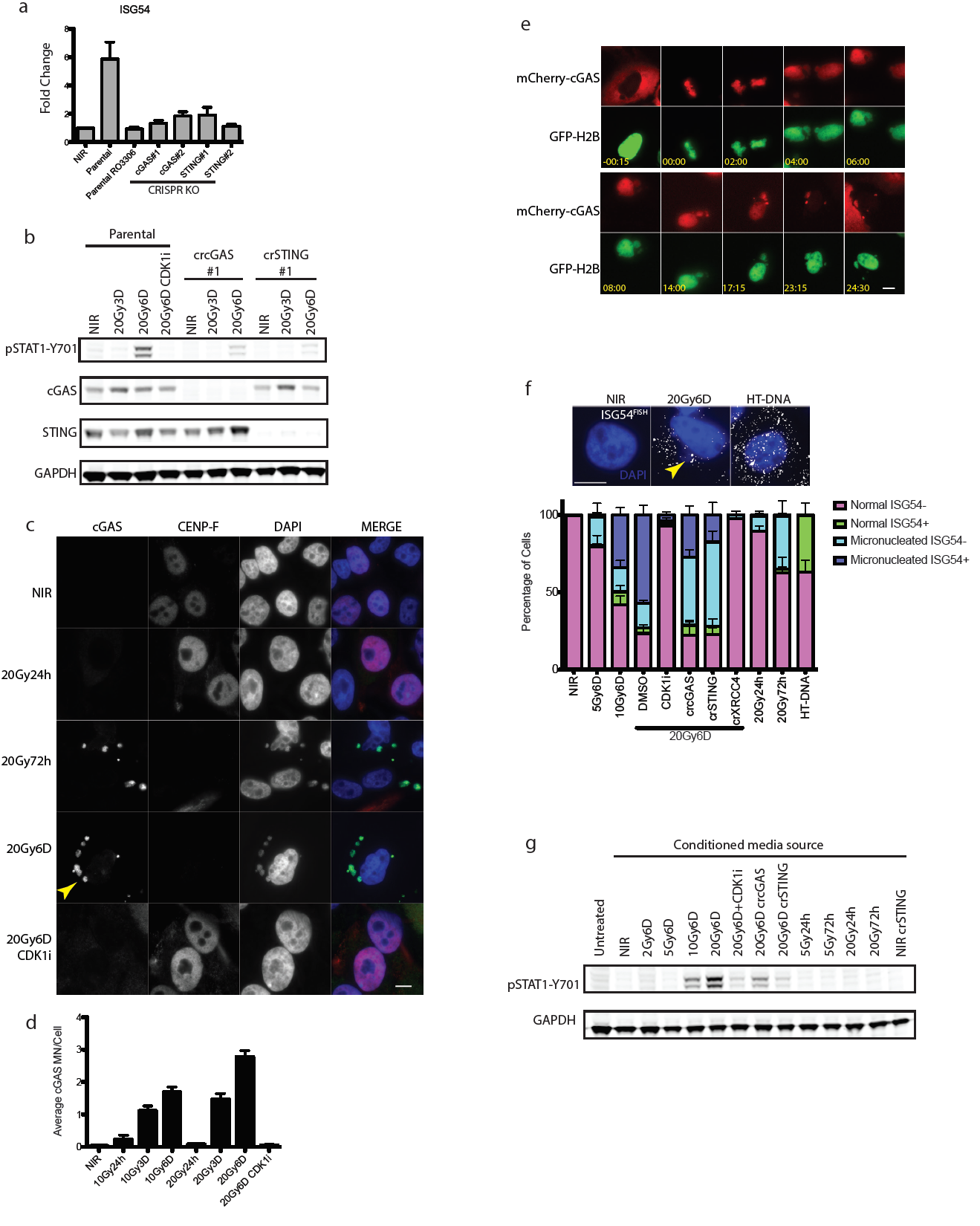
Relocalization of cGAS to micronuclei following mitotic progression triggers inflammatory signaling. **a,** RT-qPCR of ISG54 at 6 days following 20Gy in parental MCF10a cells treated +/- CDK1i (RO-3306) or derivatives harboring deletion of cGAS or STING. Error bars represent SEM of three biological replicates. **b,** Representative western blots of STAT1 activation in CRISPR-CAS9 knockout MCF10A cells for cGAS or STING. **c,** Immunofluorescence of endogenous cGAS in MCF10A cells. Arrowhead indicates a representative cGAS-positive micronucleus. Scale bar is 10µm **d,** Quantification of micronuclei in (**c**). Error bars represent SEM of three biological replicates. **e,** Extracted frames from live cell microscopy of irradiated MCF10A cells stably expressing GFP-H2B to mark chromatin and mCherry-cGAS. Inset depicts time from obvious mitotic induction in hh:mm. **f,** Representative RNA FISH images (Scale bar is 10µm) and quantification of ISG54 status as described in Methods. Arrowhead indicates an example of a micronucleus. **g,** Representative western blot of STAT1 activation of naïve cells treated with indicated conditioned media for 24h.

cGAS localization was prominent in lamin B2 positive micronuclei of CENP-F negative (G1) cells three days following IR (Fig. 3c, 3d and Extended data 3d and 3e). Synchronized cells showed cGAS localization to micronuclei concomitant with mitotic passage, G1 accumulation and STAT1 activation (Fig 2e-f and Extended data 4a). Delayed accumulation of mCherry-cGAS in micronuclei was also observed (Extended data 4b) and live-cell microscopy showed dynamic and persistent relocalization of cGAS to micronuclei following passage through mitosis (Fig 3e and Extended movie 1). Interestingly, cGAS localized to bulk chromatin during mitosis, but dissociated at completion of mitosis. cGAS concentrated in the micronuclei after a transient delay, consistent with reported changes in permeability due to delayed micronuclear envelope rupture^25^. The live cell imaging revealed ˜20% of daughter cells undergoing rapid post-mitotic apoptosis (Extended movie 2). The remaining 80% of daughter cells all contained micronuclei, the majority of which showed strong and persistent cGAS concentration (Extended data 4c).

The preceding data support a model whereby loss of micronuclear envelope integrity can provide access to damaged DNA. During migration of cells through 3µm pores (but not 8µm pores) transient DSBs are produced and the nuclear envelope is disrupted^26^. When MCF10A cells were subjected to migratory experiments we observed cGAS localization to both micronuclei and nuclear blebs (Extended data 4d and 4e). cGAS positivity of these subcellular structures still occurred in the presence of CDKi, demonstrating that mechanical nuclear rupture bypasses the need for mitotic progression. Furthermore, micronuclei generated using Aurora B kinase inhibition were cGAS positive and elicited robust STAT1 activation without irradiation (Extended data 4f and 4g).

Micronuclei harboring accumulated cGAS may act to initiate inflammatory signaling. RNA FISH revealed coincident expression of interferon-stimulated gene ISG54 specifically in cells containing micronuclei (p≤0.001 at 6D post-20Gy; Fig 3f). This effect was abrogated by knockout of cGAS, STING, or XRCC4, and by CDK1i. In an independent test of this association, expression of an IFNβ1 promoter controlled eGFP transgene increased specifically in micronucleated cells at 6 days following 10Gy (p≤0.01; Extended data 5a). These findings firmly establish a significant relationship between the presence of micronuclei and inflammatory signaling.

Only subtle induction of nuclear pIRF3 was detected by immunofluorescence in damaged cells (Extended data 5b). This suggests that chronic DNA damage induced inflammatory activation emanating from cGAS positive micronuclei may differ in magnitude from the acute signaling that occurs within hours following transfection of microgram quantities of exogenous DNA (Extended data 5c). To determine if paracrine signals were responsible for amplifying cGAS-STING mediated signaling in the setting of DSB induction, we transferred media from irradiated cells to naïve cells and assayed for STAT1 activation (Figure 3g). These experiments reveal IR induced paracrine signaling that required mitotic progression and cGAS-STING in a time and dose dependent manner.

Several case reports have described an abscopal effect defined as tumor regression outside of the irradiated field. This is most often observed in combination with immune checkpoint blockade (i.e. anti-CTLA4 antibody)^27-29^, suggesting that local radiation can be immunomodulatory. Consistent with this, we have extensively characterized how irradiation of one tumor along with immune checkpoint blockade can produce T cell-dependent responses in the contralateral unirradiated tumor using the B16 syngeneic murine model of melanoma^29,30^. To examine if mitotic progression and the cGAS-STING pathway are required for regression of unirradiated abscopal tumors after anti-CTLA4, we used this B16 model but irradiated cells *ex vivo* with or without CDK1i prior to implantation (Fig 4a). This *ex vivo* approach allowed us to eliminate the confounding impact that systemically blocking the cell cycle may have on both immune function and tumor biology.

**Figure 4:**
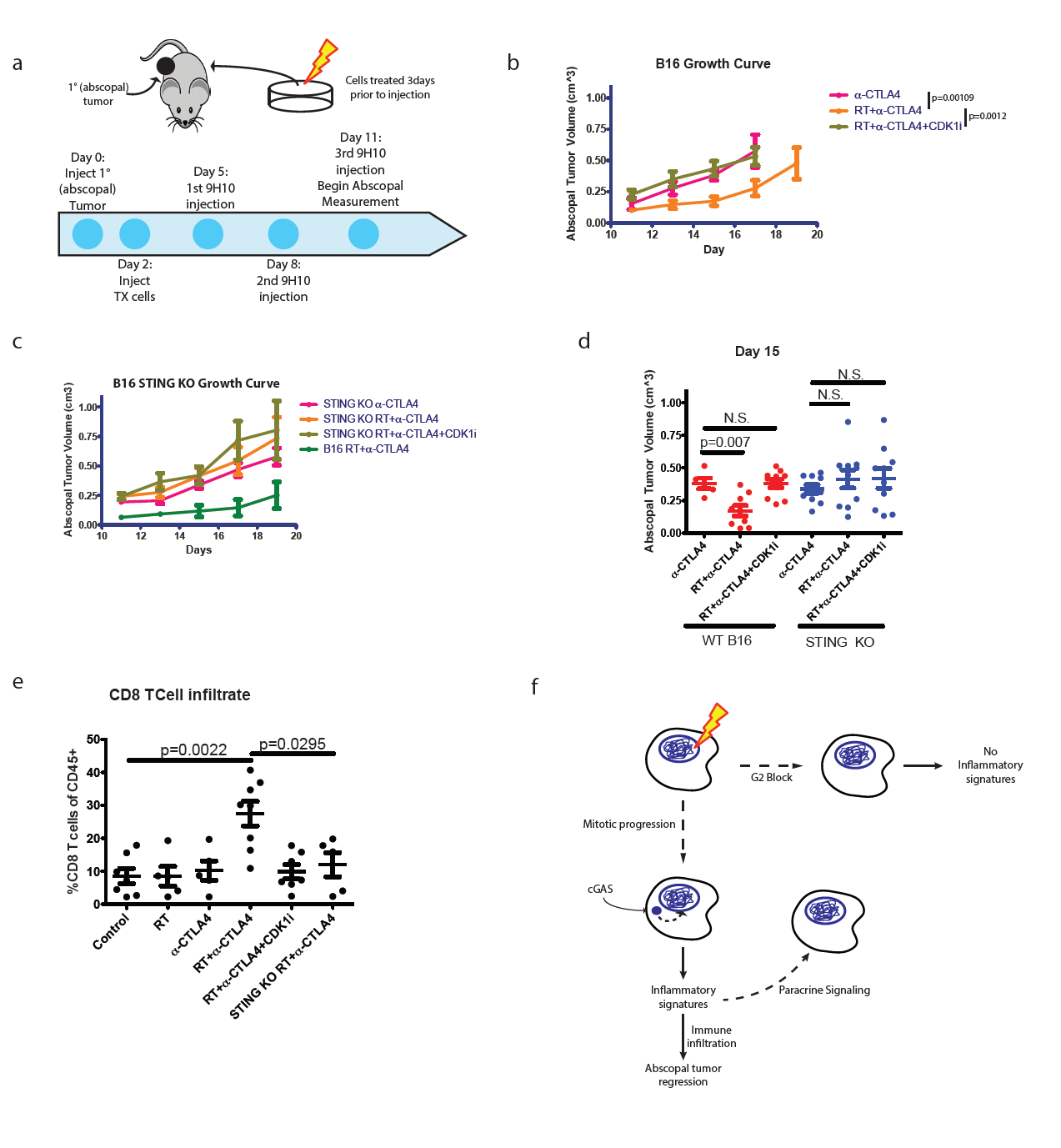
Mitotic progression and STING signaling are required for maximal anti-CTLA4 therapy driven abscopal responses in the B16 murine melanoma model. **a,** Schematic of the modified RadVax procedure. **b,** Growth of abscopal tumors following injection of untreated cells, cells treated with 10Gy 3 days before implantation, or cells irradiated in the presence of CDK1i. All mice received 9H10 anti-CTLA4 antibody as described in **a.** p-value is from the mixed effect linear model. **c,** as in (**b)** with STING knockout B16 cells. **d,** Static tumor volumes at day 15. **e,** Fraction of CD8^+^ cytotoxic T cells as a percent of CD45^+^ cells infiltrating the abscopal tumor. **f,** Model as described in the text. Pairwise comparisons by Mann-Whitney test, all error bars are SEM of biological replicates.

Irradiation of B16 melanoma cells before injection caused significant reduction in the growth of abscopal tumors after anti-CTLA4 treatment (Fig 4b). In contrast, pre-treatment with CDK1i reduced STAT1 activation and the abscopal effect (Fig 4b and Extended data 6a). Implanting irradiated B16 cells harboring STING deletion eliminated the radiation-mediated growth delay of the abscopal tumor after anti-CTLA4 irrespective of CDK1i treatment (Fig 4c and Extended data 6b). Radiation in the absence of anti-CTLA4 was insufficient to induce the abscopal effect (Extended data 6c). The abscopal tumor volume reduction was not observed when the implanted cells were treated with CDK1i or were STING deficient (Fig 4d). Loss of STING or CDK1i in the irradiated tumor also significantly reduced overall survival in the mice (Extended data 6d). A similar impact on tumor size with STING loss was noted in abscopal tumors when the contralateral tumor was irradiated directly in the mice (Extended data 6e and 6f). Consistent with a requirement for T cell responses, STING knockout or CDK1i treatment prevented the enrichment of intratumoral CD8 T cells in the abscopal tumor (Fig 4e) ^29,31^. Thus, mitotic progression with DSBs and cGAS-STING signaling is a critical component of both the intrinsic response to radiation and to host immune activation that drives regression of distal tumors.

Checkpoint adaptation and insensitivity has been described in a wide range of eukaryotic organisms^10,32-35^. Our data support a model in which imperfect cell cycle checkpoints allow passage of cells through mitosis and accumulation of micronuclei where pattern recognition occurs (Fig 4f). This represents a situation in which actively cycling cells contribute to delayed onset inflammatory signaling in the context of DSB inducing therapies. These findings suggest possibilities to modulate the host immune system and ultimately the success of genotoxic therapies.

## METHODS

### Cell lines and tissue culture

MCF10A cells were obtained from ATCC, stably transfected to express Cas9 as described below, and cultured in a 1:1 mixture of F12:DMEM media with 5% horse serum (Thermo Fisher Scientific), 20ng/mL human EGF (Peprotech), 0.5mg/ml hydrocortisone, 100ng/ml cholera toxin and 10µg/mL recombinant human insulin (Sigma). MCF10A-IPpoI cells were previously described^36^ and U2OS-IPpoI and MCF10A-AsiSI cells were prepared with identical procedures. The AsiSI cDNA was a gift of from New England Biolabs and was cloned by PCR into the pLVX-PTuner vector (Clontech) with an Estrogen receptor tag derived by PCR from the pLVX-PTuner-IPpoI vector. B16-F10 and U2OS cells were purchased from ATCC and cultured in DMEM with 10% FBS. UWB1.289 and UWB1.289+BRCA1 cells were obtained from ATCC and cultured in a 1:1 mixture of RPMI1640 and MEGM (prepared from BulletKit, Lonza) with 10% FBS added. All cells were cultured in the presence of Penicillin and Streptromycin (Thermo Fisher Scientific). Normal Prostate Epithelial Cells (PrEC) were purchased from Lonza and cultured according to recommended protocols. Cell lines were verified mycoplasma free (Lonza).

### Irradiation and cell treatments

All cells were seeded such that they reached no greater than 60-70% confluence at the time of treatment. Media was changed every 2-3 days following treatment. Irradiation was performed under ambient oxygen using a Cs-137 Gammacell irradiator (Nordion) at a dose rate of ˜0.8Gy/min. Inhibitors were added 1h before treatment and maintained until collection unless otherwise noted: DNA-PKi (NU-7441, 2µM, Tocris), CDK1i (RO-3306, 9µM, Tocris), CHK1i-1 (LY2603618, 1µM, Selleck Chemicals), CHK1i-2 (CHIR-124, 100nM, Selleck Chemicals, ATRi (VE-821, 2.5µM, Selleck Chemicals), Ruxolitinib (1µM,Cayman Chemicals), PLK1i (BI-2536, 10nM, Selleck Chemicals), Aurora Bi (AZD1152-HQPA, 10nM, Selleck Chemicals), CDK1/2/4i (R547, 0.6µM, Cayman Chemicals), CDK4/6i#1 (PD332991, 1µM, Cayman Chemicals), CDK4/6i #2 (LY28352169, 2.5µM, Cayman Chemicals) and ATMi (Ku55933, 10µM, Selleck Chemicals). PARPi (AZD-2281, 5µM, LC Laboratories) was added to UWB1.289 cells and kept on for the duration of the 6 days before harvesting. IFNβ1 (Peprotech) was added to cells at 2.5ng/mL for 4h before harvesting. 2’3’-cGAMP (InvivoGen) was added to the cells at a concentration of 10 µg/mL for 8 hours before harvesting. For nuclease induction cells were incubated in Shield-1 (1µM, Chemipharma) and 4OHT (2µM, Sigma) were added for 6h before washing twice with PBS and adding back media without drugs to allow recovery. For synchronization experiments cells were incubated in CDK1i for 16h then irradiated as necessary. Immediately following irradiation media was removed, cells washed twice in PBS and media without CDK1i was added for an appropriate duration. Herring testis DNA (Sigma) was dissolved in water and transfected into cells at the indicated concentrations using JetPrime reagent (Polyplus Transfection) according to manufacturers protocols.

### CRISPR-Cas9 knockout

Lentivirus was produced using lentiCas9-Blast (a gift from Feng Zhang, Addgene #52962)^37^ according to standard procedures, concentrated and used to infect MCF10A cells as described^36^. Cells were selected in 10µg/mL blasticidin (Invivogen) for ˜1 week and knockouts were created by infecting cells with lentivirus created using lentiGuide-Puro (a gift from Feng Zhang, Addgene #52963)^37^ followed by selection in 1µg/mL puromycin (Sigma) for another week. Cells were plated such that single colonies could be isolated after 2 weeks. Clones for NHEJ knockouts were selected by immunofluorescence staining of 53BP1 24h after 2Gy to determine cells with elevated DSB levels. Knockout clones of cGAS and STING were determined by western blotting and multiple clones were pooled for each sgRNA. sgRNA sequences are presented in Table 1.

**TABLE 1:**
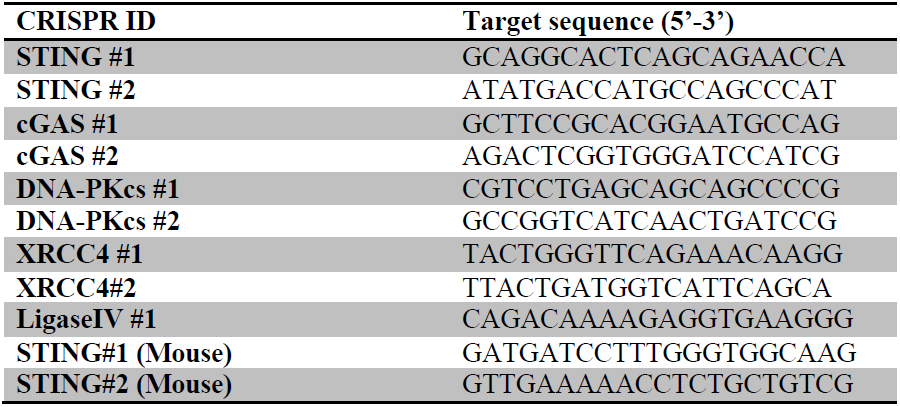
CRISPR target sequences

B16-F10 STING knockouts were generated as previously described^30^, sgRNA sequences are presented in Table 1. Single cell clones were screened for knockout by adding cGAMP as described below and monitoring ISG induction. Four confirmed knockouts from each guide were combined together to create the B16-F10 STING knockout cell line.

### Western blotting

Western blotting was performed using standard methods. Briefly, cells were scraped in PBS and lysed in RIPA buffer (150mM NaCl, 1% NP-40 alternative, 0.5% Sodium deoxycholate, 0.1% SDS, 40mM Tris pH8.0) with 1mM Benzamidine HCl, 1µg/mL antipain, 5µg/mL aprotinin, 1µg/mL leupeptin, 1mM Phenylmethansulfonyl fluoride, 10mM NaF, 2mM imidazole, 1.15mM Sodium molybdate, 4mM Sodium tartrate and 2mM Sodium orthovanadate (Sigma)) for 20 minutes on ice and cleared by centrifugation. Protein was quantified using Bradford reagent and 20-30µg of protein were run per lane of 4-12% Bis-Tris gels (Thermo Fisher Scientific) using either MOPS or MES buffer. Protein was transferred to nitrocellulose membrane at 350mA for 2h in transfer buffer (15% methanol, 191mM glycine, 25mM Tris base, 0.1% SDS, pH8.3). Membranes were blocked in 5% non-fat milk powder in PBS+0.2% Tween 20 (PBST) and incubated overnight at 4°C in primary antibody diluted in blocking solution. Primary antibodies are listed in Table 2. After extensive washing in PBST blots were incubated for 1h at room temperature (RT) in Licor blocking solution (1:1 with PBST) containing IRDye 800CWDonkey anti-Rabbit or IRDye 680LT Donkey anti-Mouse secondary antibody (1:10000,Licor) as required. Blot were scanned on a Licor Odyssey scanner at a medium resolution setting and exported to ImageJ (NIH).

**TABLE 2:** Antibodies and dilutions used

| **Target** | **Company** | **Catalog#/Clone** | **Dilution** |
| --- | --- | --- | --- |
| **pSTAT1-Y701** | Cell Signaling | 9167/58D6 | WB: 1/1000 |
| **STAT1 total** | Cell Signaling | 9176/9H2 | WB: 1/1000 |
| **pATM-S1981** | Abcam | Ab81292/EP1890Y | WB: 1/1000 |
| **DNA-PKcs** | Thermo Fisher Scientific | MA5-13238/18-2 | WB: 1/1000 |
| **LigaseIV** | Abcam | Ab193353/EPR16531 | WB: 1/1000 |
| **XRCC4** | BD Biosciences | 611506/Clone 4 | WB: 1/1000 |
| **GAPDH** | Cell Signaling | 2118/14C10 | WB: 1/5000 |
| **γH2AX-S139** | Millipore | 05-636/JBW301 | IF: 1/2000 |
| **53BP1** | Novus Biologicals | NB100-904 | IF: 1/1000 |
| **pH3-Ser10** | Millipore | 06-570 | WB: 1/1000 |
| **CENP-F** | BD Biosciences | 610768/Clone 11 | IF: 1/800 |
| **cGAS** | Cell Signaling | 15103/D1D3G | IF: 1/500<br>WB: 1/1000 |
| **STING** | R&D Systems | MAB7169 | WB: 1/1000 |
| **IFITM1** | Proteintech | 11727-3-AP | WB: 1/1000 |
| **pIRF3-Ser396** | Cell Signaling | 29047/D601M | IF: 1/50 |

### RT-qPCR

RNA was isolated using TRIZOL reagent under standard conditions. Reverse transcription and quantitative PCR was carried out as previously described^36^. Primers are listed in Table 3.

**TABLE 3:**
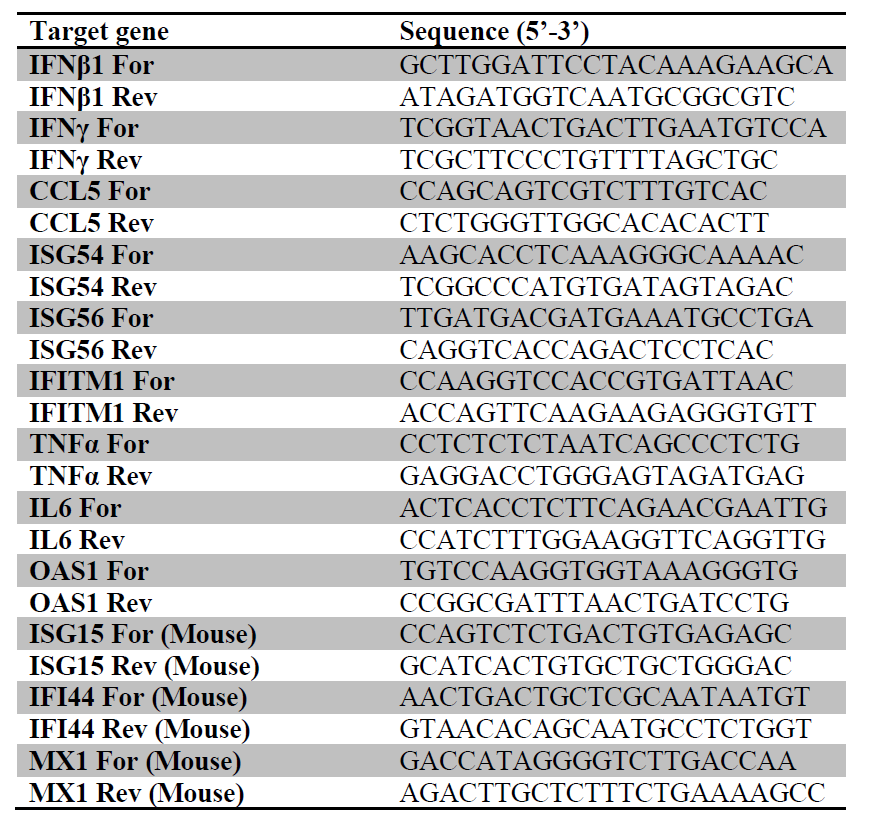
Primers for qPCR

### Immunofluorescence

Cells were seeded onto coverslips and following treatment were processed as described^36^ with the following exceptions. Coverslips were mounted in Vectashield Hard Set mountant containing DAPI (Vector Labs). For pIRF3 staining Tween20 was omitted from all buffers. Images were captured using a Coolsnap Myo camera (Photometrics) connected to a Nikon Eclips 80i microscope using a 60X oil immersion or 20X air objective and Nikon NIS-Elements software. Images were prepared for publication using FIJI (NIH) and all channels of a given figure were adjusted equivalently only for brightness and contrast. Antibodies are listed in Table 2. Micronucleated cells were classified manually by distinct staining by DAPI of structures outside of the main nucleus. To enumerate cGAS positive micronuclei these structures were counted manually for each field and expressed as a percentage of total cells within the field. Total nuclear intensity was calculated on a per cell basis using FIJI (NIH). Graphs and statistics were prepared using Prism Graphpad.

### Flow cytometry

Cells were harvested from a single well of a 6-well plate using trypsin and washed 1x in PBS before fixation with dropwise addition of ice-cold 70% ethanol. After fixation on ice (or storage at −20°C) cells were washed 1x in PBS and permeabilized in 0.25% Triton X-100/PBS solution on ice for 15 minutes. Cells were washed 1x in PBS and resuspended in 1% bovine serum albumin/PBS solution containing 1µl/sample of Phospho-histone H3 (Serine 10) antibody conjugated to Alexa Fluor 488 (DC28, Cell Signaling). After washing 1x in PBS cells were resuspended in a solution of PBS containing 50µg/mL Propidium iodide (Santa Cruz Biotechnology) and 100µg/mL RNAse A (Roche). Flow cytometry was performed on a FACSCalibur (BD Biosciences) and analysed using FlowJo software. Single cells and G1/S/G2 peaks were manually gated. The gates for phospho-H3 positive cells were chosen by comparison to a population in which the antibody was omitted during processing.

### Plasmid construction

The plasmid backbone was isolated from pLVX-pTUNER-N using Q5 polymerase and primers (5’-ggatccGGTATTTAAATAGATCCTCTAGTAGAGTCGGTGTC-3’, 5’-accggtgcggccgcCGATCCCGCCCCTCTCCC -3’) for Gibson assembly. The mCherry fragement was isolated from the pHFUW-mCherry-TRF-FOKI plasmid^38^ with Q5 polymerase and primers (5’-agaggatctatttaaataccggatccATGGTGAGCAAGGGCGAG-3’, 5’-ccatcttcatctcgagCTTGTACAGCTCGTCCATGC-3’). These two fragments were joined by Gibson assembly (New England Biolabs) and digested with XhoI/NotI (New England Biolabs). cGAS was amplified by PCR from Human Lung Fibroblast DNA (Biosettia) using primers: 5’-catctcgagATGCAGCCTTGGCACGGAAAG-3’ and 5’-atggcggccgcTCAAAATTCATCAAAAACTGGAAACTCATTGTTTC-3’. After digestion with XhoI/NotI the fragment was inserted using a Quick Ligation Kit (New England Biolabs). MCF10A cells were infected with virus as described above.

### Live cell microscopy

The pBABE-H2B-GFP plasmid was a gift from Fred Dick (Addgene plasmid #26790)^39^ and retrovirus was created using standard protocols and introduced into MCF10A cells expressing mCherry-cGAS. Dual mCherry/GFP positive cells were isolated by cell sorting and seeded onto 35mm Mattek dishes (Mattek). Cells were synchronized with CDK1i and released according to the procedure described above. Imaging was started immediately after release. Imaging was carried out with a Deltavision microscope enclosed in a 37°C humidified chamber equilibrated with 5% CO_2_. Images were captured using a Photometrics CoolSnapHQ camera using a 40X air objective. Single plane images were acquired every 15 minutes for 20-24h. Focus was stable throughout with minor manual adjustments over the imaging timecourse. This procedure was repeated three times during which ˜100 daughter nuclei could be analysed, corresponding to 50 mitotic events. The mobility of the cells limited the ability to follow all mitotic events to completion. H2B-GFP bodies separate from the main nuclei were identified as micronuclei and apoptotic cells were identified by rapid and severe H2B-GFP condensation before complete dissociation and loss from the culture surface (Supplementary movie 2).

### Senescence associated β-Galactosidase assay

Cells were seeded at ˜2x10^5 cells per well of a 6-well plate, treated 24h after seeding with IR and the indicated agents, and 6 days later were fixed and stained for SA-βGal as described ^40^. Three technical replicates were performed within at least 2 independent biological experiments.

### Conditioned media experiments

Cells for all treatments were plated at a constant density and irradiated as described. Conditioned media was obtained by changing to fresh media 24h before collection and filtering though a 0.22µm filter. This conditioned media was transferred to naïve cells and protein lysates for western blotting were collected 24h later.

### RNA FISH

RNA FISH was performed as described with modifications^36^. Cells were fixed on coverslips with 2% formaldehyde in PBS and permeabilized at least 24h and up to 1 week at 4°C in 70% ethanol. After a brief rinse in RNA Wash buffer (10% formamide/2X SSC) cells were inverted into hybridization buffer containing Custom Stellaris® FISH Probes designed against ISG54 (NM_001547) by utilizing the Stellaris® RNA FISH Probe Designer (Biosearch Technologies, Inc., Petaluma, CA) available online at www.biosearchtech.com/stellarisdesigner. After overnight hybridization cells were washed twice for 30 minutes at room temperature in RNA wash buffer, once for 2 minutes in 2X SSC and mounted in vectashield. Imaging was performed using a 60x objective as described above. Using ImageJ, all images from a given treatment were combined in a montage and background subtracted with a rolling ball radius of 10. A threshold was chosen based on HT-DNA transfected cells to eliminate any remaining background that was not a clear diffraction limited RNA FISH spot. Cells with at least 10 FISH spots were counted as positive for ISG54.

### IFNβ–GFP Reporter assay

The IFNβ-luciferase reporter plasmid was a kind give of R. Vance^41^. Using Gibson assembly the luciferase reporter gene was swapped with eGFP (from pBABE-H2B described above) to form the IFNβ-GFP cassette. A fragment derived from the pLKO.1-Hygro plasmid (a gift from B. Weinberg, Addgene #24150) that omitted the U6 promoter sequence was combined with the IFNβ-GFP cassette to generate the pLKO.1-Hygro-IFNβ-GFP reporter. A stable MCF10A cell line was derived as described above by viral infection and selection with 100ug/mL hygromycin (Invivogen). Activation of GFP was monitored by microcopy in fixed cells. The threshold used for quantification of GFP positive cells was kept constant throughout all experiments.

### Transwell migration

For migration through transwells (Corning Inc.), cells were seeded at 300,000 cells/cm^2^ onto the topside of the filter membrane and left to migrate in normal culture condition for either 9 or 24 hours. For experiments involved CDK1i, cells were exposed to 9 µM CDK1i and DMSO for 1 hour prior to the seeding onto Transwells. Transwell membrane was fixed in 4% formaldehyde (Sigma) for 15 minutes, followed by permeabilization by 0.25% Triton-X (Sigma) for 10 minutes, blocked by 5% BSA (Sigma) and overnight incubation in various primary antibodies: lamin-A/C (Santa Cruz), lamin-B (Santa Cruz) and cGas (Cell Signalling). Finally, the primary antibodies were tagged with the corresponding secondary antibodies for 1.5 hours (ThermoFisher). DNA was stained with 8µM Hoechst 33342 (ThermoFisher) for 15 minutes. Confocal imaging was performed on a Leica TCS SP8 system using a 63x/1.4 NA oil-immersion lens.

### Modified RadVax procedure

Five to seven week old female C57BL/6 mice were obtained from Charles River and maintained under pathogen free conditions. Mice were divided randomly into cages upon arrival and are randomly injected and measured. All animal experiments were performed according to protocols approved by the Institute of Animal Care and Use Committee of the University of Pennsylvania. On the day before the experiment was to begin B16-F10 cells were treated as necessary *in vitro* with 10Gy of IR with or without CDK1i. As necessary the CDK1i was added 1h before irradiation and maintained until cell isolation and injection on day 2 below. On Day 0 untreated B16-F10 cells (5x10^4^) in 50µL of PBS were mixed with an equal volume of Matrigel (BD Biosciences) and injected into the right flank. On Day 2 5x10^5^ B16-F10 parental or STING knockout cells treated as described above were mixed with matrigel and injected on the opposite flank. On Days 5, 8 and 11 anti-CTLA4 antibody (9H10; BioXCell) was administered interperitoneally at 200µg per mouse. Volumes were measured using calipers starting at day 11 and calculated using the formula L X W^2^ X 0.52, where L is the longest dimension and W is perpendicular to L. Overall survival and flow cytometry for tumor infiltrating CD8^+^ T-cells was performed as described^30^. For the *in situ* irradiation experiments of Extended Data 4e and 4f experiments were performed using the Small Animal Radiation Research Platform (SAARP) as previously described^29^.

## Acknowledgements

We thank members of the Greenberg lab for critical discussion and A. Jackson (University of Edinburgh, Scotland) for sharing unpublished results. We are also grateful to R. Vance (UC Berkeley) for providing the IFN-β promoter driven plasmid that we subsequently modified to drive eGFP expression. This work was supported by NIH grants CA17494, GM101149, and CA138835 (to RAG), and by funds from the Basser Center for BRCA

## Author Contributions

SMH and RAG designed the study and wrote the manuscript. SMH performed most of the experiments. JLB designed and performed in vivo experiments described in Figs 4 and S6. JI designed and performed experiments described in Fig S5a-b. AJM and DED provided guidance for the design of experiments and edited the manuscript. The authors declare no competing financial interests.

## FIGURE LEGENDS

**Figure S1:**
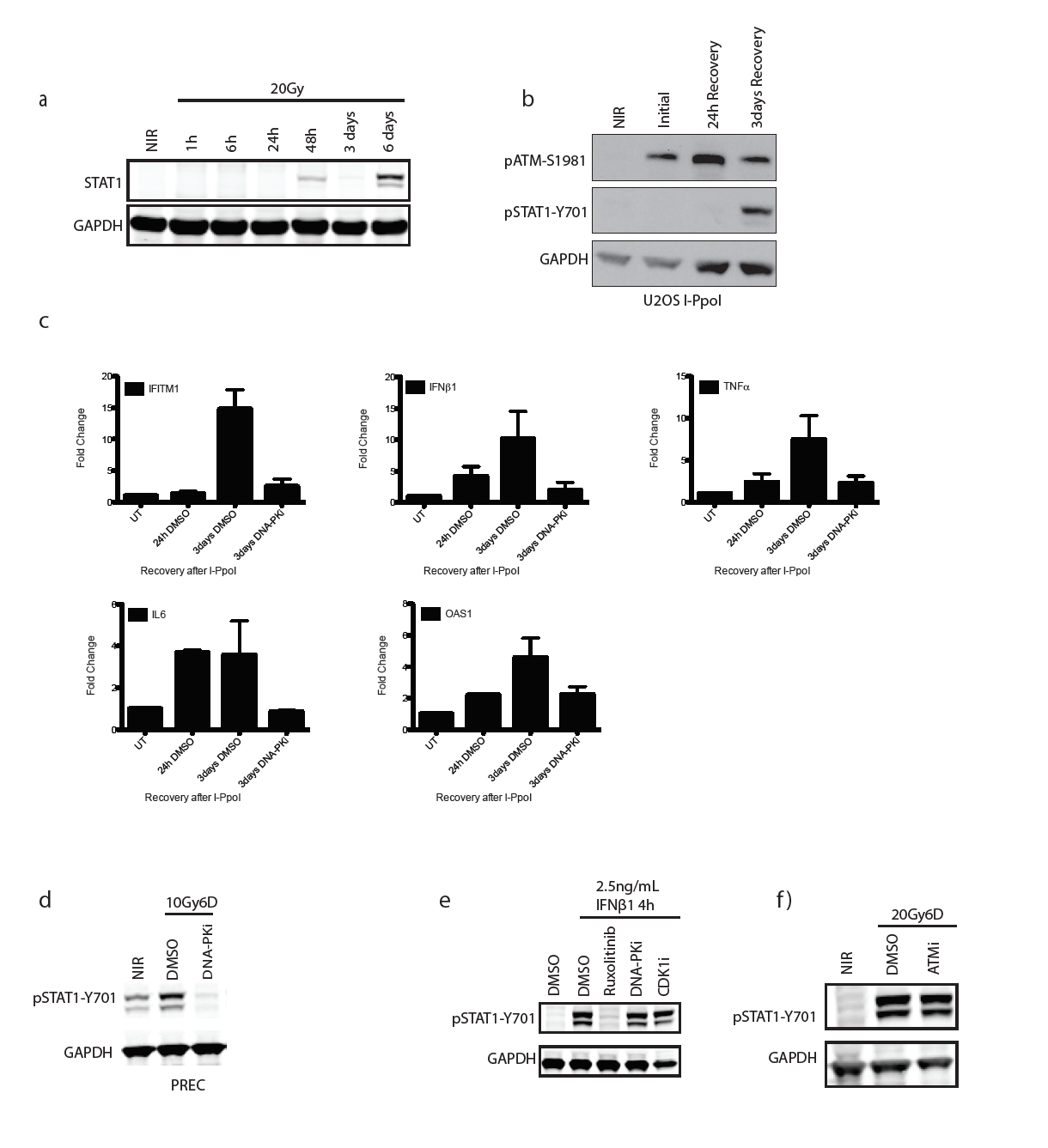
**a,** Total STAT1 protein levels are increased in a time-dependent manner after IR. **b,** U2OS cells were monitored for STAT1 activation during recovery from I-PpoI nuclease. **c,** Gene expression changes were monitored for indicated genes during recovery from I-PpoI damage in MCF10A cells (see figure 1c). **d,** Prostate epithelial cells (PREC) were analyzed for STAT1 phosphorylation at 6 days following 10Gy. **e,** STAT1 phosphorylation following IFNβ1 treatment is not DNA-PK or CDK1 dependent. **f,** pSTAT1 activation measured by western blot is unchanged by ATMi.

**Figure S2:**
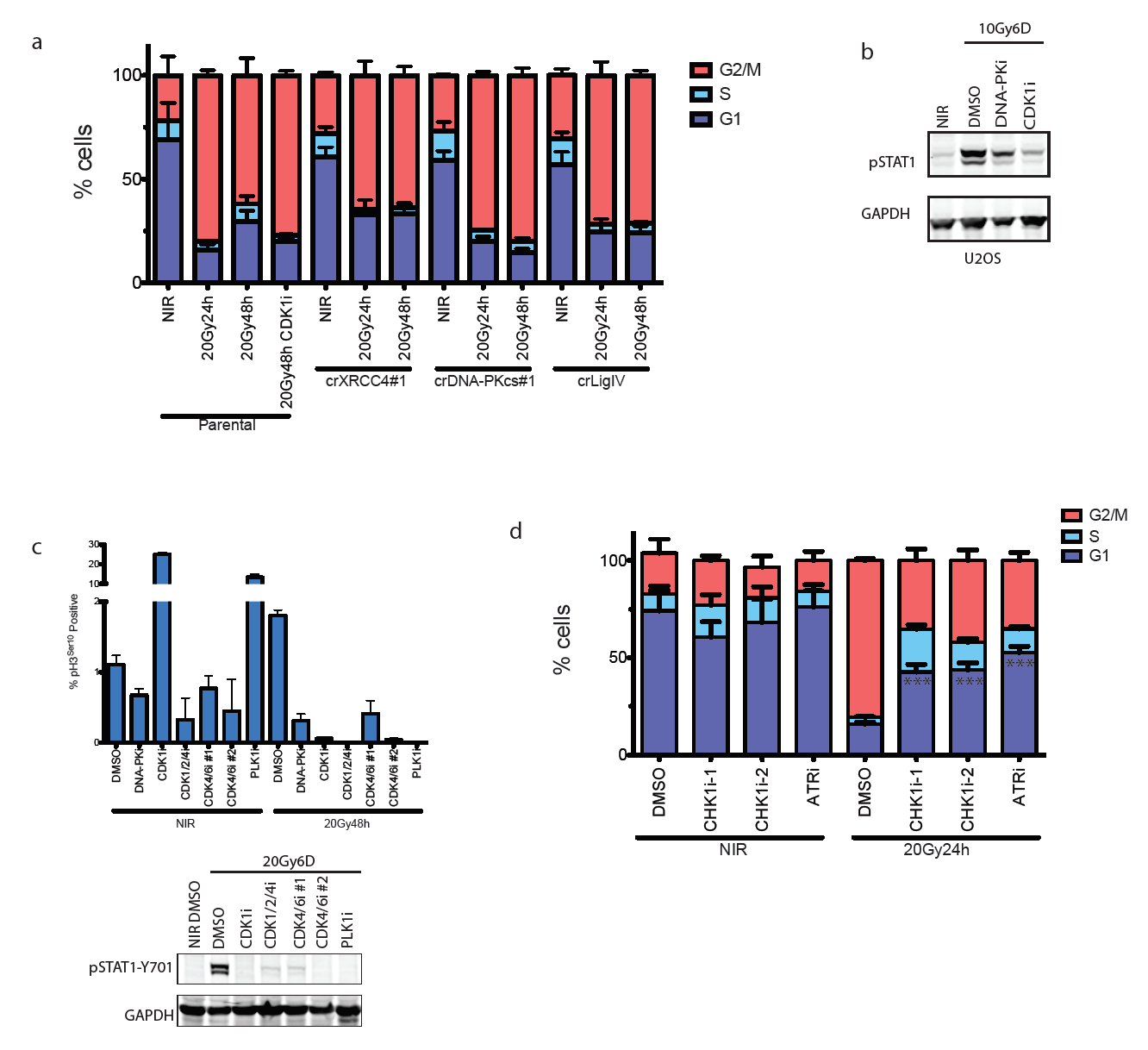
**a,** Cell cycle profiles of MCF10A cells were monitored by flow cytometry of Propidium iodide-stained cells treated as indicated. **b,** U2OS cells were monitored for STAT1 activation in the presence of the indicated inhibitors. **c,** H3 (Ser10) phosphorylation in MCF10A cells was measured by flow cytometry and expressed as a percentage of total single cells. Error bars represent SEM of at least 2 biological replicates. Western blots show loss of STAT1 activation in treatments corresponding to the flow cytometry data. **d,** Cell cycle profiles of PI-stained MCF10A cells treated as indicated. G1 populations of irradiated, inhibitor treated cells are significantly higher than of vehicle (DMSO) control. ***p<0.0001 with Dunnett’s multiple comparison.

**Figure S3:**
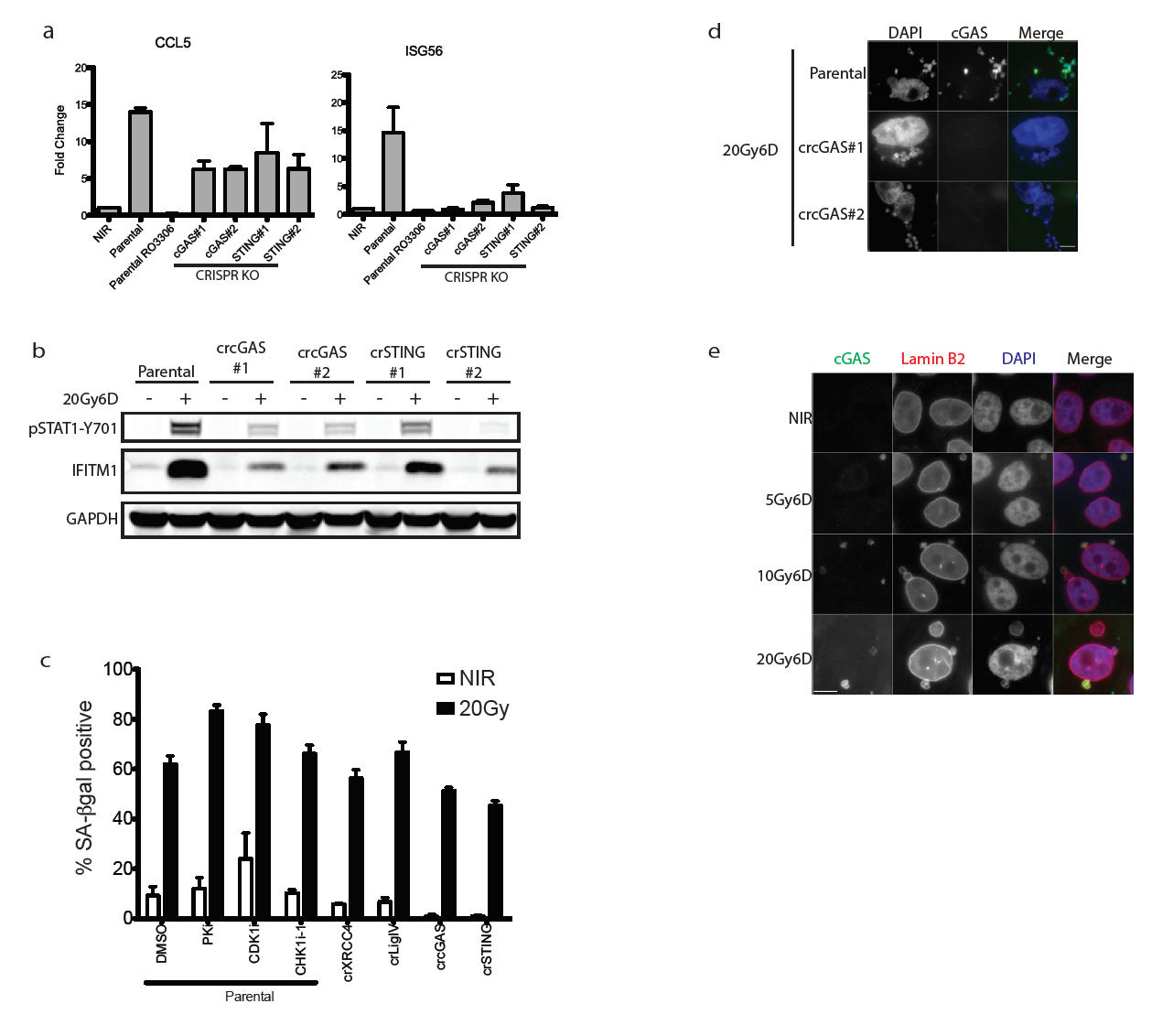
**a,** RT-qPCR of CCL5 and ISG56 at 6 days following 20Gy in parental MCF10A cells treated +/- CDK1i (RO-3306) or derivatives harboring deletion of cGAS or STING. Error bars represent SEM of three biological replicates. **b,** Knockouts for two separate CRISPR-Cas9 sgRNA for cGAS or STING cause similar reductions in STAT1 signaling. **c,** Senesence-associated β-galactosidase staining of 6D following indicated treatments. Error bars represent SEM of at least 2 biological replicates. **d,** Representative immunofluorescent staining of cGAS shows loss of staining in cGAS knockout cells. Scale bar is 10um. **e,** Immunofluorescent staining shows costaining of Lamin B2 in cGAS positive micronuclei. Scale bar is 10µm.

**Figure S4:**
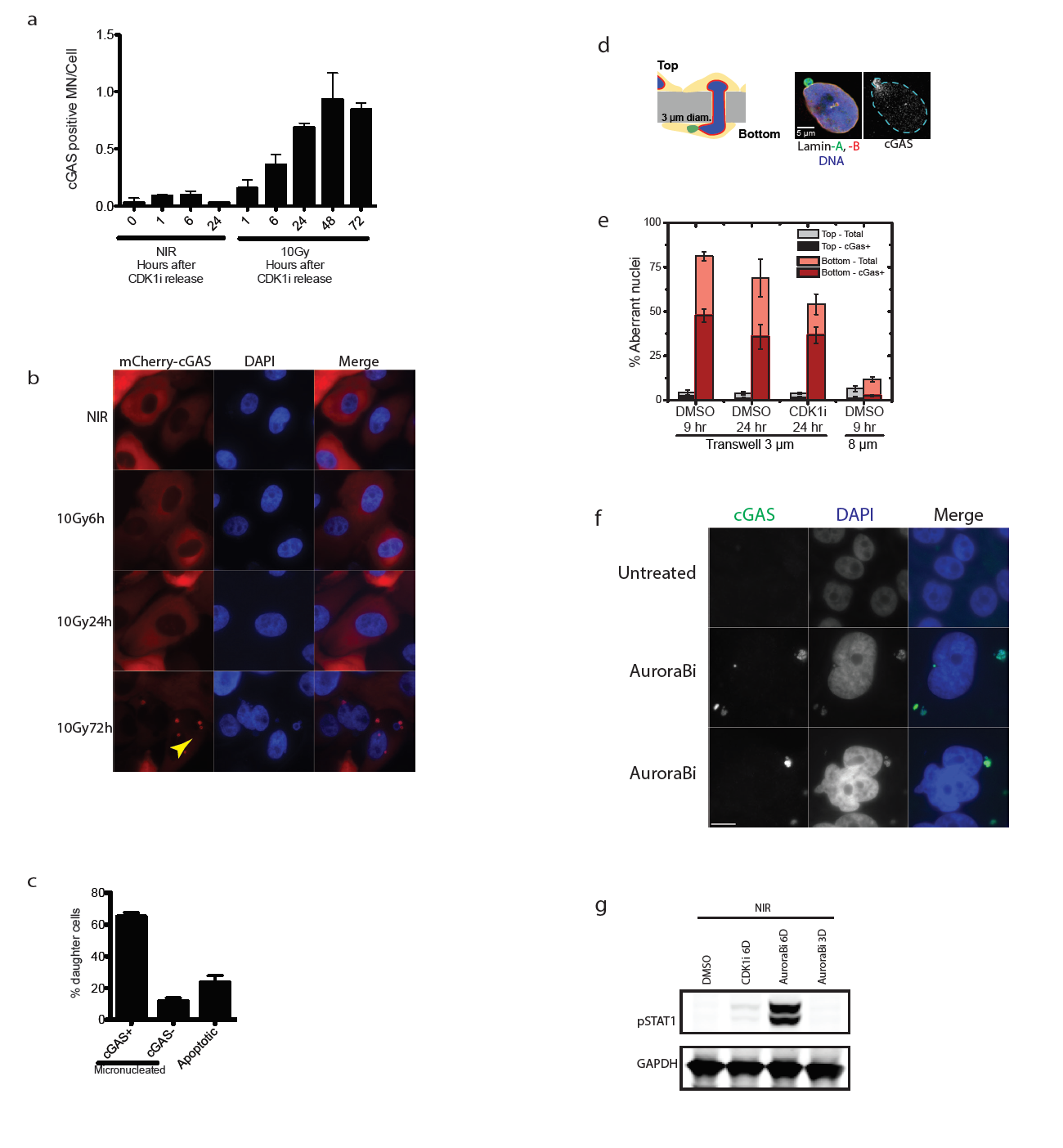
**a,** Cells in figure **2d** were analyzed for the fraction of cGAS positive micronuclei following release from CDK1i. **b,** mCherry-cGAS expressing cells were monitored by IF following 10Gy. Arrowhead indicates mCherry-cGAS positive micronucleus. **c,** Histogram represents the fraction of daughter cells with cGAS positive micronuclei or that underwent apoptosis during live-cell imaging. All non-apoptotic daughter cells were micronucleated after division. Error bars are SEM of three biological replicates (n=99 total daughters). **d,** Schematic of nuclear migration transwell system. Blue is DAPI, Green is Lamin A and Red is Lamin B. cGAS is shown in grayscale. **e,** Quantification of aberrant nuclei (nuclear blebs and micronuclei) that are cGAS positive in the transwell migration assay. **f,** Immunofluorescent staining in an untreated and two representative cells after 6 day treatment with Aurora B kinase inhibition. Scale bar is 10um and similar patterns were observed in two independent experiments. **g,** Representative western blot of STAT1 activation in non-irradiated cells treated with Aurora B inhibitor for 3 or 6 days.

**Figure S5:**
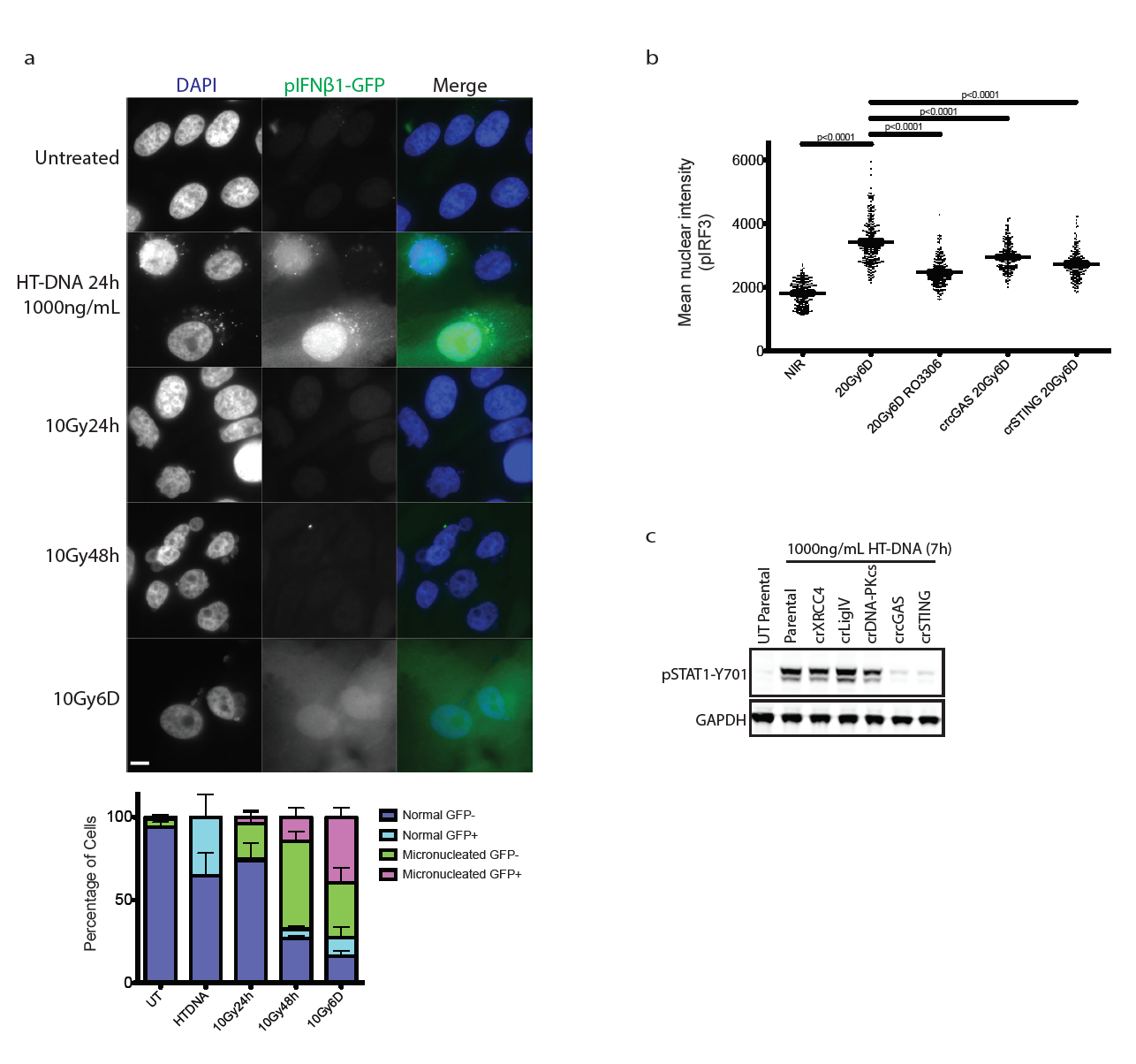
**a,** Representative images of pIFNβ-GFP reporter cells treated as indicated. Scale bar is 10µm. Quantification is as described in Methods and error bars represent SEM of 2 biological replicates. **b,** Mean nuclear intensity of pIRF3 staining was quantified. p-values are based on pooled data from 3 independent experiments and calculated by 1-way ANOVA. **c,** Representative western blot of STAT1 activation following transfection of herring testis DNA (HT-DNA) in indicated CRISPR-Cas9 knockout MCF10A cell lines shows STAT1 activation in DNA-PK and all NHEJ deficient cells, but not in cGAS STING knockouts.

**Figure S6:**
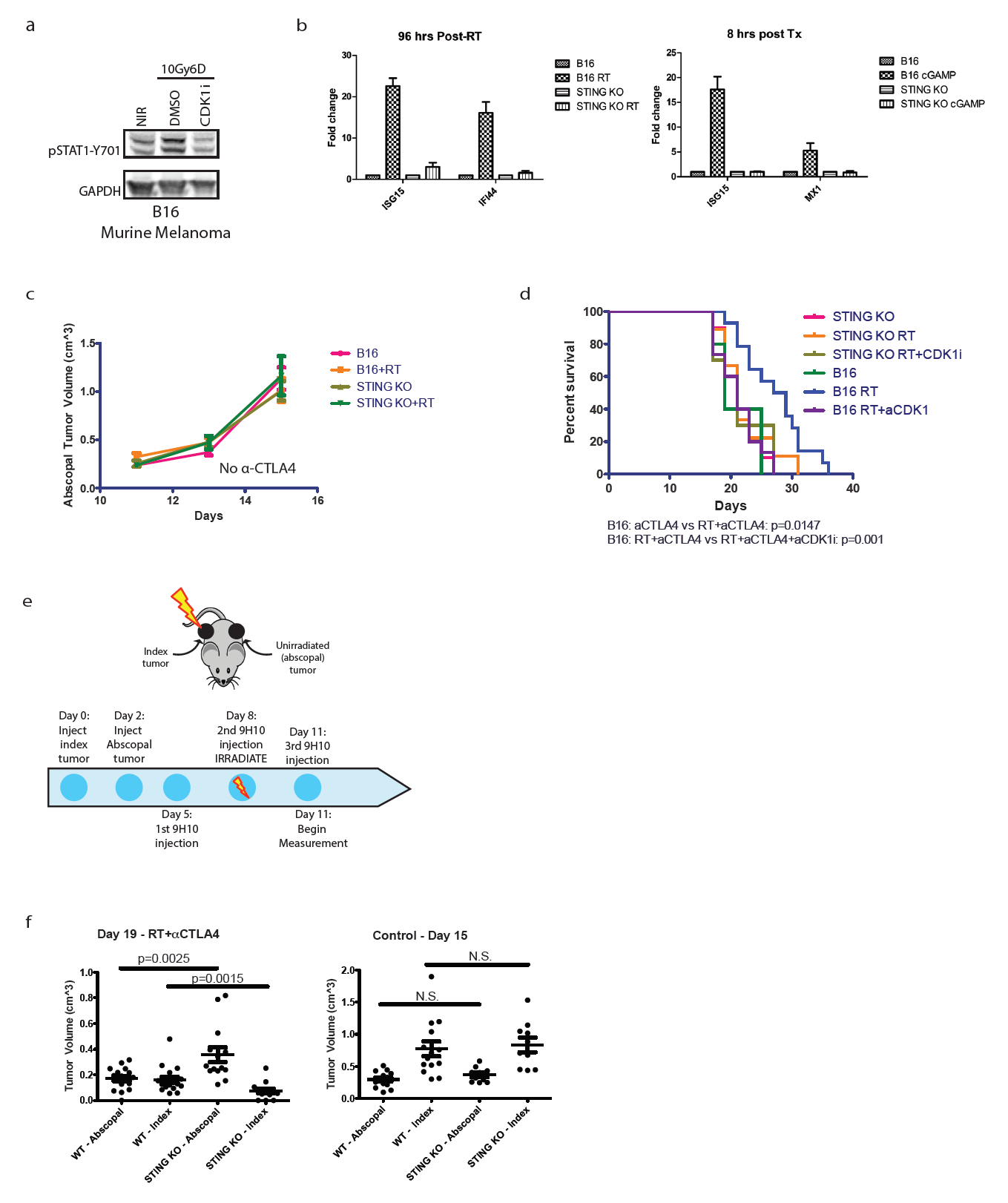
**a,** Representative western blot of STAT1 activation in B16 cells with and without CDK1i. **b,** RT- and cGAMP-induced gene induction is absent in STING knockout B16 cells. **c,** Injection of B16 parental or STING knockout cells (+/- RT) without combination anti-CTLA4 treatment is insufficient to induce an abscopal effect. **d,** Overall survival of mice when B16 parental or STING knockout were injected after indicated treatment. All mice received anti-CTLA4 antibody. P-value calculated by log-rank test. **e,** B16 tumors are injected into opposite flanks of mice and treated as indicated. The index tumor is irradiated with 20Gy and both the index and abscopal tumors are measured starting on day 11 post-injection. **f,** irradiation of the index tumor leads to an abscopal response that is dependent on STING (left panel). This response is not seen in unirradiated mice (right panel).

**Supplementary movie 1:** Representative movie of mitotic cell becoming micronucleated shown in Fig 3e. H2B-GFP (green) and mCherry-cGAS (red) were imaged as described in the text. The inset timestamp is in hh:mm and rendered at 1 frame per second.

**Supplementary movie 2:** Representative movie of post-mitotic apoptotic cell as quantified in Fig 3e. H2B-GFP (green) and mCherry-cGAS (red) were imaged as described in the text. The inset timestamp is in hh:mm and rendered at 1 frame per second.

